# RTCpredictor: Identification of Read-Through Chimeric RNAs from RNA Sequencing Data

**DOI:** 10.1101/2023.02.02.526869

**Authors:** Sandeep Singh, Xinrui Shi, Syed Basil Ahmad, Tommy Manley, Claire Piczak, Christopher Phung, Yunan Sun, Sarah Lynch, Aadi Sharma, Hui Li

## Abstract

Read-through chimeric RNAs are gaining attention in cancer and other research fields, yet current tools often fail in predicting them. We have thus developed the first read-through chimeric RNA specific prediction method, RTCpredictor, utilizing a fast ripgrep algorithm to search for all possible exon-exon combinations of parental gene pairs. Compared with other ten popular tools, RTCpredictor achieved top performance on both simulated and real datasets. We randomly selected up to 30 candidate read-through chimeras predicted from each software method and experimentally validated a total of 109 read-throughs and on this set, RTCpredictor outperformed all the other methods. In addition, RTCpredictor (https://github.com/sandybioteck/RTCpredictor) has less memory requirements and faster execution time.

## INTRODUCTION

Identification of chimeric RNAs generated by intergenic splicing has opened up a new paradigm of transcriptional diversity wherein many events have been discovered and experimentally validated in multiple studies (1–8). Among them, a subset of chimeric RNAs have been identified as diagnostic markers and therapeutic targets in cancer patients (9–11) and many others are being explored for their potential (8, 12–14). Chimeric RNAs have also been extensively reported in normal healthy tissues and cells (15–17) thereby challenging the traditional view as being unique to neoplasia. A plethora of databases hosting chimeric RNAs have been made and extensive annotations have been provided to give insights about their functional relevance (3, 18–21). Based on the coordinates from where the break points occur, chimeric RNAs can be grouped into three different types: (i) Read-through, (ii) inter-chromosomal, and (iii) intra-chromosomal.

Read-through chimeric RNAs, also known as products of cis-Splicing of Adjacent Genes (cis-SAGe) are generated when RNA polymerase extends its transcription beyond a single gene to include a nearby gene on the same strand, and exons from these two parental genes are spliced together. Although read-through transcripts have been widely considered as a class of chimeric RNA (22), some categorize them as potential regular transcripts (13, 23, 24). In the past, they were generally assumed to be biologically insignificant for tumorigenesis (25–27), because of the absence of genomic rearrangements. Therefore, they were often considered as false positive events and were discarded by experimental biologists while screening for cancer chimeras (28). The same assumption was made by many computational biologists in designing their prediction methods, and these events were filtered out using distance-based criteria (29). However, a number of read-through chimeric RNAs with clear clinical and biological relevance have been reported. Rickman *et al*. (30) identified chimeric RNA SLC45A3-ELK4 in prostate cancer and demonstrated its biomarker capabilities and Zhang *et al*. (31) classified it as a cis-SAGe event by cracking down its mechanism of formation. Zhang *et al*. also demonstrated the correlation of this read-through chimera with Gleason score, which is not seen with parental gene transcripts (31). Varley *et al*. (32) screened 168 breast samples to identify and experimentally validate *SCNN1A-TNFRSF1A* and *CTSD-IFITM10* read-throughs present in breast cancer cell lines and primary tumors but not detected in normal tissues. By performing analysis of deep RNA-Seq in prostate cancer and reference samples, Nacu *et al*. (33) identified *MSMB-NCOA4* read-through which may play some functional roles in cancer. Recently, we data mined multiple cancer types from TCGA and explored the landscape of chimeric RNAs in multiple cancer types. We identified and experimentally validated several read-through chimeric RNAs (*LHX6-NDUFA8* in cervical cancer (34), *D2HGDH-GAL3ST2* in prostate cancer (35), *BCL2L2-PABPN1* in bladder cancer (36), *RRM2-C2orf48* in colorectal cancer (37)) and showed their potential to be used as diagnostic markers and therapeutic targets. Impressively, same clinical correlation was often not seen with the parental gene transcripts, and sometimes even opposite trend was observed as in the case of *RRM2-C2orf48* (37). In addition, several papers have reported physiological read-through chimeric RNAs that are necessary in normal cells/tissues (16, 38–40). These data support that read-through chimeric RNAs represent a novel class of transcriptional products, and some of them may play important roles in physiology and tumorigenesis, or serve as ideal biomarkers or therapeutic targets.

Clearly, many more read-through chimeric RNAs are still hidden in the unmined data and it is the need of the hour to pay attention to them and to include them in the chimeric RNA analysis pipeline. The main limitation in the high-throughput analysis of read-throughs is the lack of computational methods specifically designed for predicting them. Although a general chimeric RNA prediction software can be used to predict these events, they lack the sensitivity as demonstrated by the latest benchmark study of sixteen chimeric RNA prediction methods on read-through class of chimeric RNAs (29). Moreover, many of these software methods apply stringent criteria where most of these events, if not all, are filtered out (29). In this study, we describe a prediction method, Read-Through Chimeric RNA Predictor (RTCpredictor), specifically designed for the identification of read-through chimeric RNAs. We have used a simulated and real dataset of read-throughs, and showed its superior performance on these datasets. Apart from high sensitivity, our method is fast and scalable in high-throughput settings.

## MATERIALS AND METHODS

### Datasets

To demonstrate the performance of our prediction method, we have used a simulated dataset and three real datasets having 50, 46, 6 and 11 read-throughs respectively. The detail of datasets is given below:

ChimPipe simulated dataset: It consists of 250 chimeric RNAs with 50 read-throughs among them. Briefly, the dataset was created using ChimSim module of ChimPipe package (41) wherein 250 simulated chimeric RNAs were mixed with 102,149 Gencode (version 19) transcripts including the parental genes of the chimeric RNAs. Three paired-end libraries were created with read length of 101, 76 and 50 bp and roughly total number of reads as 31, 42 and 64 million respectively. Consistent with the read-through criteria defined in our previous benchmark study where the distance between 5’ and 3’ breakpoint coordinates should be within ≤70 kb, we excluded 14 out of 50 read-throughs having distance >70 kb and showed performance of our method and other methods on 36 simulated read-throughs. The list of these 36 chimeric RNAs along with their breakpoint coordinates is given in Supplementary Table S1.

Qin *et al*. real dataset: It consists of 62 chimeric RNAs with 46 read-throughs among them. These read-throughs were identified from LNCaP cell lines and were also experimentally validated by RT-PCR and sanger sequencing (42). Consistent with our previous benchmark study (29), for this study we used two runs (SRR1657556 and SRR1657557 from SRA study ID SRP050061) and applied quality filtering with NGSQC Toolkit (43) with default parameters to select high quality reads. The read length of both runs is 101 with roughly total number of reads after filtering as 62 and 58 million respectively. Just like in simulated dataset, here too, we excluded two out of 46 read-throughs having distance between 5’ and 3’ breakpoint coordinates >70 kb. We also excluded nine read-throughs whose 5’ and 3’ breakpoint coordinates did not map to the end and start of the participating exons respectively. Therefore, we evaluated the performance of our method and other methods on 35 real read-through. The list of these 35 chimeric RNAs along with their breakpoint coordinates is given in Supplementary Table S2.

TCGA bladder dataset: It contains six experimentally validated read-through chimeric RNAs which were identified from TCGA bladder cancer RNA-Seq data using EricScript software (44). Since each read-through was detected in multiple samples, we randomly selected three samples for each read-through and ran all the software methods on these three samples. If the software method detected the read-through in any of the three samples, we then considered that the software successfully re-identified the read-through. EricScript software was not compared on this dataset because it is the tool that was used to identify these six read-throughs in the TCGA bladder cancer study. The sample ids corresponding to each read-through is provided in Supplementary Table S3.

TCGA colorectal dataset: It contains eleven experimentally validated read-through chimeric RNAs which were identified from TCGA colorectal cancer RNA-Seq data using EricScript software (37). We used same approach as in the TCGA bladder cancer dataset i.e. we randomly selected three samples for each read-through and ran all the software methods (except EricScript). We considered the software successfully re-identified the read-through if it detected the read-through in any of the three samples. The sample ids corresponding to each read-through is provided in Supplementary Table S4.

### Performance parameters

We have used Sensitivity, Positive Predictive Value (PPV) and F-measure to calculate the performance of our prediction method on simulated and real datasets. Briefly, Sensitivity is the percentage of correctly predicted true positive events; PPV is fraction of true positive events among all positive predicted events and F-measure (also known as F1-score or F-score) is a balanced score which captures both sensitivity and PPV. These parameters are standard parameters used in benchmarking studies and the formula for calculating these parameters can be accessed from previous studies (29, 45). Although, we have reported all the three parameters mentioned above, it should be noted that PPV and F-measure are not well suited for comparing the performance of chimeric RNA prediction methods on real datasets. This is because we don’t know the exact number of true chimeric RNAs present in real datasets and the PPV/F-measure of a software will drop if it predicts an unknown but true chimeric RNA (29). Moreover, PPV and F-measure parameters are also not appropriate to be used in the present study because our software method predicts only read-through class of chimeric RNAs while other software methods predict all classes of chimeric RNAs; therefore, only sensitivity can be used for a one-to-one comparison. All previous benchmark studies have compared the performance of methods based on gene pairs but in this study, we showed the performance of our method and other methods based on breakpoint coordinates i.e. at the isoform level. Prediction at isoform level is more difficult and thus more stringent than predicting at gene pair level.

### Computational time and memory calculation

To calculate time and memory load required for running this prediction method, we used ‘time’ command which is built-in command in linux operating system. We reported time in minutes and memory load in GB and consistent with our previous benchmark study (29) we used same high-performance computing system and SRR1657556 sample to test computational load so that one-to-one comparison with other methods can be done. Briefly, a single core on a single node having Intel Xeon processor (E5-2630 @2.40GHz) was used to run individual software methods.

### Experimental validation

RNA extracted from the LNCaP cell line was reverse transcribed via Verso cDNA synthesis Kit (Thermo Scientific) (46). Primer pairs were designed using Primer3 targeting the relative upstream and downstream exons of the junction sequence. The primer design follows the rules: 1) Primer length is between 18bp to 20bp. 2) Primer Tm is between 57°C to 63°C. 3) Primer GC% is between 40% to 60%. 4) Poly-X of primer is no more than 3. 5) Primer targeting site is >50bp away from the junction site whenever possible. Primer sequences are listed in Supplementary Table S5. A Step One Plus Real-Time PCR System (Applied Biosystems) was used to perform SYBR Green-based qRT-PCR experiments. Amplified products were separated using agarose gel electrophoresis. Proper size product bands were purified using the PureLink Quick Gel Extraction Kit (Invitrogen) and submitted for Sanger sequencing (Genewiz). To validate the chimera candidates, PCR product sequences obtained from Sanger sequencing were aligned to the human genome through the BLAT tool.

## RESULTS

### RTCpredictor algorithm

We designed the algorithm to conduct exhaustive search of junction sequences obtained by all possible exon combinations of neighboring genes against RNA-Seq data. We consider potential read-throughs as those events where both 5’ and 3’ parental genes are transcribed in the same direction and the distance between 5’ and 3’ breakpoints is ≤70 kb, which is consistent with the criteria used in our previous benchmark paper (29). We extracted 20 bp sequence from each side of the breakpoints and combined them to form the junction sequence of 40 bp in length. Using hg19 genome and Ensembl version 75 annotation file, 4,967,029 exon combinations representing 9,781,826 unique sequences (junction sequences and their reverse complement) were extracted from 110,877 gene pairs with distance within ≤70 kb (Figure 1). Using ripgrep string search software (https://github.com/BurntSushi/ripgrep), these 9.78 million junction sequences, each corresponding to a unique read-through transcript, were searched against RNA-Seq data to identify read-through events. Since this approach does not require mapping RNA-Seq data to the genome, the algorithm is very fast and free of errors introduced by mapping step.

**Figure 1.**
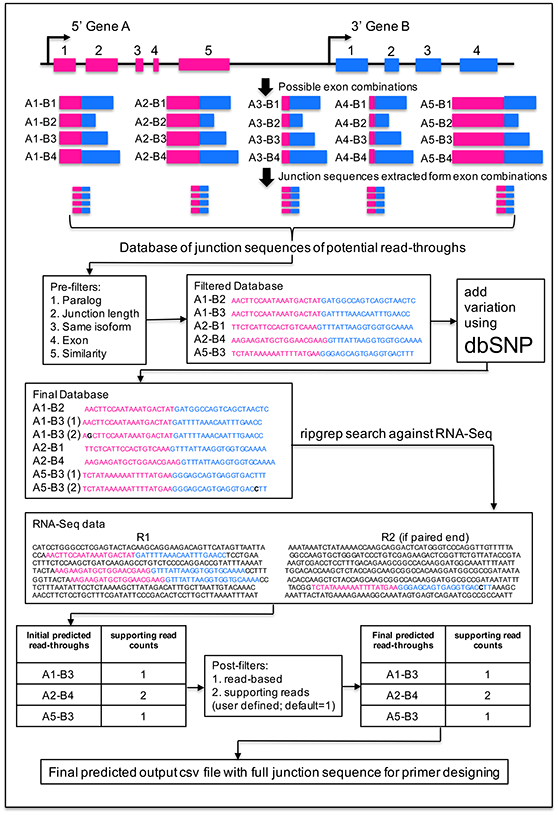
Flowchart of RTCpredictor method. Exemplary genes A and B are located on the same strand with five and four exons respectively. There are 20 possible exon combinations, given that the distance between breakpoints is <=70KB, which is used to generate a database of junction sequences of potential read-throughs. Pre-filters are applied to discard potential false positive events. dbSNP is used to map and add common variants on junction sequences to generate the final database. This final database is used to search against RNA-Seq input data using ripgrep for the initial predicted read-throughs. Post-filters are applied to exclude some more potentially false positive events to generate the final list of predicted read-throughs. The annotations such as genomic coordinates and junction sequences are included in the output file which will help the users to design primers for their experimental validation.

### Filters to discard potential false positive events

We implemented several filters to identify and discard potential false positive events. First five of these filters are implemented before the step involving ripgrep search against RNA-Seq data i.e. they discard potentially false positive junction sequences (exon combinations) (Figure 1). The rest of the two filters are implemented after ripgrep search. The filters are: (1) Paralog filter: paralog genes were downloaded from ensemble version 75. If 5’ and 3’ genes of the potential read-through are paralogs of each other, those combinations were discarded (9,303,622 junction sequences). (2) Junction sequence length filter: If 5’ or 3’ exon length is <20 bp then junction sequence of length 40 bp cannot be constructed; therefore, those combinations were discarded (9,146,174 junction sequences). (3) Same isoform filter: If 3’ exon of the junction sequence is also present in 5’ gene and it is next to the 5’ exon and vice versa, then it is discarded (9,002,938 junction sequences). These events arise from overlapping genes. (4) Exon filter: The junction sequence is discarded if it matches to any exon in the hg19 genome. A blast database of all the exon sequences of the hg19 genome was constructed and blast search of junction sequences against exon database was performed. Sequence identity >=90% and alignment length covering >=90% of the junction sequence was used as criteria to discard them (8,999,838 junction sequences). (5) Similarity filter: If 5’ exon is similar to any exon of the participating 3’ gene and vice versa, these events were discarded. A blast search of 5’ exon (or 3’ exon) sequence was performed against all exons of 3’ gene (or 5’ gene) and sequence identity ≥70% with alignment length coverage of ≥70% was used to discard these events (8,789,234 junction sequences). After these filters, a database file was constructed containing 8,789,234 junction sequences with their breakpoint coordinates, ready to be used for ripgrep against RNA-Seq data. (6) read-based filter: After ripgrep search, number of supporting reads were calculated for each read-through. If any supporting read was identified to support more than one read-through then that read was discarded and the supporting read counts were recalculated. If the discarded read was the only read supporting any read-through then that candidate read-through entry was discarded. (7) number of supporting reads: This filter is an optional filter provided to the users to select read-throughs with minimum number of supporting reads.

### Mapping variation in junction sequences

Since RTCpredictor searches for exact match of junction sequences against RNA-Seq data, it may miss read-throughs having variation in the junction sequence in different individuals. To tackle this issue, we identified SNPs with common allele frequency (CAF) ≥10% mapping on the junction sequences and added new junction sequences reflecting common variations in our database file. We downloaded SNP data from dbSNP (version b151) using GRCh37p13 genome and used bedtools (version 2.27.1) (47) to intersect with our junction sequences and identified 9,372 such SNPs.

To enhance the applicability of RTCpredictor across genomes of different organisms, we have also provided a script to make the database files for the genome of interest and provided the instructions on how to download the prerequisite input files to run the script. Further, the distance parameter between 5’ and 3’ breakpoints can be modified by the user with 70,000 bp set as default value.

### Performance on ChimPipe simulated dataset

Our method RTCpredictor achieved 94.4%, 94.4% and 83.3% sensitivity on ChimPipe PE101, PE76 and PE50 datasets respectively, if only read-based filter is applied (Figure 2a and Supplementary Table S6). However, it also gave lot of predictions (10032, 9995 and 9322 on PE101, PE76 and PE50 respectively), many of which can be potentially false positives. Although RTCpredictor predicted all 36 read-throughs on PE101 and PE76 datasets, *CT45A2-CT45A3* and *HBA2-HBA1* were discarded by read-based filter and thereby sensitivity dropped to 94.4%. On PE50 dataset, four additional read-throughs (*AIMP2-ANKRD61*, *ALOX15B-AC129492.6*, *ZFYVE28-MXD4* and *ZNF837-ZNF497*) were missed by RTCpredictor due to short read length (50bp) of PE50 dataset since the junction sequences searched by RTCpredictor are 40bp in length. After adding the paralog filter, four additional read-throughs were discarded and the sensitivity dropped to 83.3%, 83.3% and 72.2% on PE101, PE76 and PE50 datasets. Importantly, adding four more filters (Junction sequence length filter, Same isoform filter, Exon filter and Similarity filter) did not affected sensitivity, however, it drastically reduced the total number of predictions (213, 210 and 191 on PE101, PE76 and PE50 respectively) by discarding potentially false positive events (Figure 2b and Supplementary Table S6). The total number of predicted read-throughs remained the same, even with the addition of common SNP variations (CAF≥10%).

**Figure 2.**
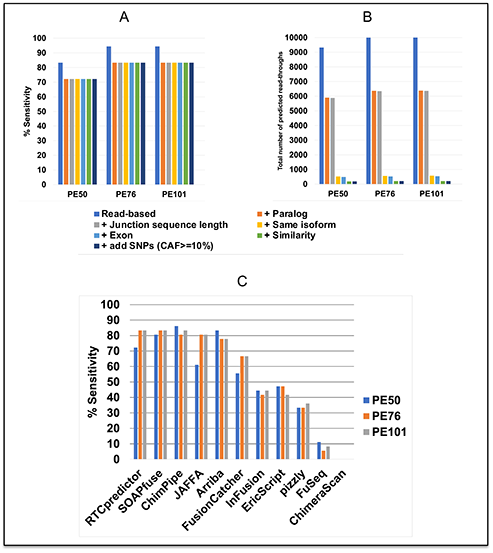
Performance of RTCpredictor and its comparison with other methods on ChimPipe simulated (PE50, PE76 and PE101) datasets. (a) % Sensitivity achieved by RTCpredictor and (b) total number of predicted read-throughs, after sequential addition of different filters. (c) Comparison of % Sensitivity of RTCpredictor with other methods.

We also compared the performance of other methods with RTCpredictor on ChimPipe simulated dataset. In consistent with our previous benchmark study (29), we chose the following ten most popular chimeric RNA prediction methods: (i) JAFFA (23), (ii) SOAPfuse (48), (iii) EricScript (49), (iv) FusionCatcher (50), (v) ChimPipe (41), (vi) pizzly (51), (vii) InFusion (52), (viii) Arriba (26, 27), (ix) FuSeq (53) and (x) ChimeraScan (54). We did not include INTEGRATE, MapSplice, STARChip, STAR-Fusion and TopHat-Fusion methods since their sensitivity on all read-through datasets was less than 5% (29). We also excluded ChimeRScope method since it does not provide the breakpoint coordinates of predicted chimeric RNAs. For all these software methods, wherever applicable, the parameters were adjusted to include read-through chimeric RNAs in their prediction (Supplementary Table S7). We downloaded the output files of all the software methods benchmarked in our previous study for calculating their performance (29).

Based on sensitivity, on ChimPipe PE101 dataset, our method RTCpredictor along with SOAPfuse and ChimPipe were the top performers (83.3%) while JAFFA (80.6%) and Arriba (77.8%) secured 2^nd^ and 3^rd^ spot (Figure 2c and Supplementary Table S8). On ChimPipe PE76 dataset as well, RTCpredictor along with SOAPfuse were the top performers (83.3%) while ChimPipe and JAFFA collectively secured 2^nd^ spot (80.6%) and Arriba (77.8%) scored the 3^rd^. On ChimPipe PE50 dataset, the performance of the methods was in the following order: ChimPipe (86.1%) > Arriba (83.3%) > SOAPfuse (80.6%) > RTCpredictor (72.2%) > JAFFA (61.1%) (Figure 2c). The drop in the performance of RTCpredictor on PE50 dataset is not surprising. In fact the performance is still impressive as RTCpredictor searches junction sequences of length 40bp against short reads of length 50bp. Therefore, it is expected that some chimeric RNAs may be missed. Based on PPV, on ChimPipe PE101 and PE50 dataset, ChimPipe (0.15, 0.15) performed the best followed by RTCpredictor (0.141, 0.136) and SOAPfuse (0.138, 0.13); and on ChimPipe PE76 dataset, RTCpredictor (0.143) performed the best followed by ChimPipe (0.139) and SOAPfuse (0.135) (Supplementary Table S8). The performance based on F-measure followed the same trend as PPV (Supplementary Table S8).

### Performance on Qin *et al*, TCGA bladder and TCGA colorectal cancer real datasets

RTCpredictor achieved 97.1% sensitivity on Qin *et al* real dataset missing only one read-through (*BRCA1-VAT1*). However, a total of 9,474 read-throughs were predicted. Paralog filter discarded an additional read-through (*AKAP8L-AKAP8*) dropping the sensitivity to 94.3%. After adding four more filters, the sensitivity remained the same, and the total number of predictions drastically dropped to 841. After the addition of common SNP variations (CAF≥10%), nine more read-throughs were predicted, making the total to 850 (Figure 3a,b and Supplementary Table S6).

**Figure 3.**
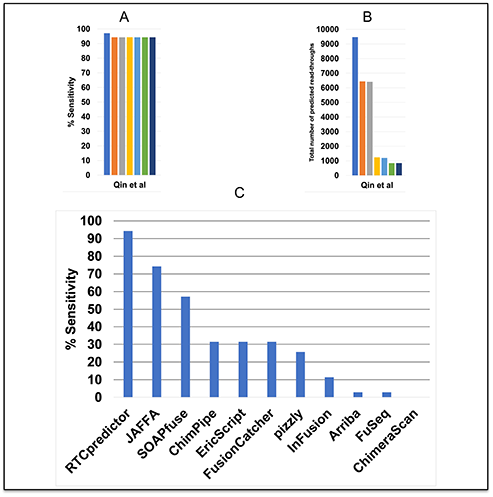
Performance of RTCpredictor and it’s comparison with other methods on Qin *et al*. real dataset. (a) % Sensitivity achieved by RTCpredictor, (b) total number of predicted read-throughs, after sequential addition of different filters, and (c) Comparison of % Sensitivity of RTCpredictor with other methods.

On TCGA bladder cancer dataset, RTCpredictor re-identified three out of six known read-throughs which are ACKR2-KRBOX1, CHCHD10-VPREB3, and SLC2A11-MIF (Supplementary Table S9). On TCGA colorectal cancer dataset, RTCpredictor uncovered five out of eleven known read-throughs which are CTSD-IFITM10, GCSH-C16orf46, METTL21B-TSFM, SLC2A11-MIF, and TRIM2-MND1 (Supplementary Table S10).

We also compared the performance of other methods with RTCpredictor on all real datasets. On Qin *et al* real dataset, based on sensitivity, RTCpredictor achieved significantly higher sensitivity (20% more) as compared to the second-best method JAFFA. RTCpredictor (94.3%) was the top performer while JAFFA (74s.3%) and SOAPfuse (57.1) secured 2^nd^ and 3^rd^ spot respectively (Figure 3c and Supplementary Table S8). Based on PPV, the performance of the methods was in the following order: SOAPfuse (0.233) > ChimPipe (0.172) > InFusion (0.105) > FusionCatcher (0.099) > RTCpredictor (0.039) (Supplementary Table S8). The performance based on F-measure followed similar trend with RTCpredictor (0.075) at 5^th^ spot (Supplementary Table S8).

On TCGA bladder cancer dataset, out of six known read-throughs, RTCpredictor re-identified three read-throughs (50% sensitivity) followed by SOAPfuse, FusionCatcher and ChimPipe which re-identified two read-throughs (33.3% sensitivity) each, while FuSeq and pizzly re-identified a single read-through (16.7% sensitivity). JAFFA, Arriba, ChimeraScan and InFusion were not able to identify any read-through (0% sensitivity) (Supplementary Table S9). On TCGA colorectal cancer dataset, RTCpredictor re-identified six read-throughs (54.5% sensitivity) followed by SOAPfuse, and ChimPipe which re-identified three read-throughs each (27.3% sensitivity) while JAFFA and FuSeq re-identified two read-throughs (18.2% sensitivity) followed by FusionCatcher and InFusion which re-identified a single read-through (9.1% sensitivity). Arriba, ChimeraScan and pizzly were not able to identify any read-through (0% sensitivity) (Supplementary Table S10).

### RTCpredictor uncovered more read-throughs in ChimPipe simulated dataset

The authors of the ChimPipe dataset simulated only 250 chimeric RNAs out of which 50 were read-throughs, however, RTCpredictor predicted 213, 210 and 191 on PE101, PE76 and PE50 respectively. Moreover, all 210 read-throughs from PE76 are also predicted in PE101 and all 191 read-throughs from PE50 are also predicted in PE76. Since sequences from transcriptome were added by the authors in ChimPipe simulated dataset, and the transcriptome itself has some transcripts annotated as “read-through”, we reasoned that some of our predicted read-throughs could match these annotated read-throughs from the transcriptome. In Ensembl version 75 annotation from hg19 genome, 371 transcripts from 110 genes are annotated as read-through. We performed a blast search of our predicted read-through sequences (extracted from combining 5’ and 3’ exon partners) against these 371 transcript sequences to identify if any of our predicted read-throughs matches with annotated read-throughs. We used ≥90% sequence identity along with ≥90% alignment length to identify such hits. 27, 27 and 23 predicted read-throughs from PE101, PE76 and PE50 matched with the annotated read-throughs from the transcriptome. Therefore, these are additional ones other than 50 read-throughs which were simulated in the ChimPipe data by the authors. It is also possible that other predicted read-throughs may be read-throughs of the transcriptome but they are not still annotated.

### Experimental validation of additional read-throughs on Qin *et al.* dataset

We believe that Qin *et al*. real dataset may have more than 46 true read-through events. To search for these additional true events hidden in the data, we randomly selected up to 30 read-through chimeric RNAs from the predicted list of each software method and performed experimental validation. We followed the following steps to select candidates for each software method: (i) The chimeric RNAs with 5’ and 3’ genes on the same chromosome and strand and having the distance between their breakpoints ≤ 70000 bp, were selected as potential read-throughs. (ii) We removed 46 events from this list that were already experimentally validated by the authors of Qin *et al*. study. (iii) We also removed 40 known read-throughs which we have previously published within the list of GTEx chimeric RNAs and are present across all 53 human tissues. (iv) We removed read-throughs having clone-based gene names starting with letters “RP”. After performing the above steps, we randomly selected 30 chimeric RNAs for software methods which produced more than 30 events in their list, and all the candidates for tools that predicted less than 30 events. We got 208 unique read-through chimeric RNAs after combining the selected list from all software methods. Finally, we performed experimental validation of these 208 read-throughs and successfully validated 109 of them using RT-PCR and sanger sequencing by manually confirming that the sequence matches the junction sequence of the read-through chimeric RNA. We call these 109 experimentally validated additional read-throughs as an extended Qin *et al* real dataset. The list of 208 read-throughs along with their validation status is given in Supplementary Table S5. RT-PCR bands, sanger sequence results and its visualization on genome browser, for five experimentally validated read-through chimeric RNAs from extended Qin *et al* real dataset and predicted by RTCpredictor are represented in Figure 4.

**Figure 4.**
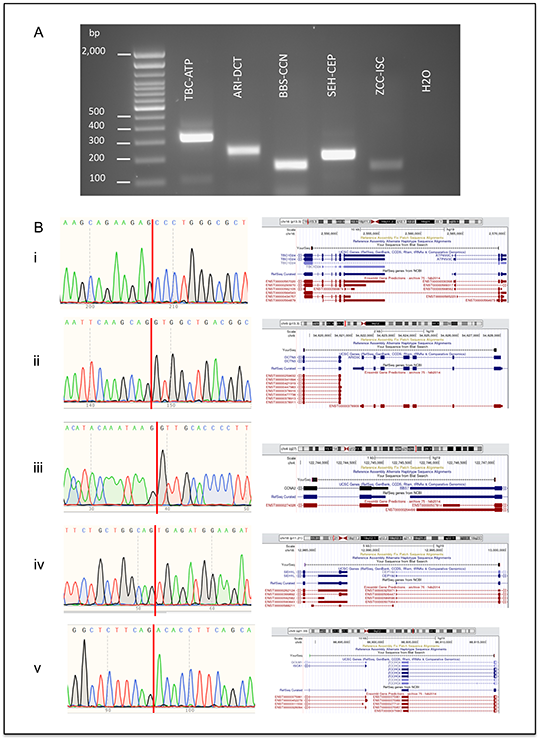
Experimental validation results of a subset of predicted read-through chimeric RNAs from extended Qin *et al*. real dataset. (a) Experimental validation using RT-PCR. Bands at the bottom of the gel are primer dimers. (b) Junction sequences obtained from sanger sequencing validation and its visualization on UCSC genome browser for the following exemplary read-through chimeric RNAs (i) *TBC1D24-ATP6V0C*, (ii) *ARID3C-DCTN3*, (iii) *BBS7-CCNA2*, (iv) *SEH1L-CEP192* and (v) *ZCCHC6-ISCA1*.

Performance comparison of all the methods on extended Qin *et al* real dataset

With 109 experimentally validated read-throughs from this study, to the best of our knowledge, Qin *et al* dataset now hosts the largest number of experimentally validated read-through chimeric RNAs. We then calculated the performance of all the methods on this real dataset. RTCpredictor performed the best with 57.8% sensitivity followed by JAFFA (42.2%) and pizzly (22.9%) while Arriba (0.9%), InFusion (0.9%) and ChimPipe (6.4%) performed poorly (Supplementary Table S11). After adding 35 read-throughs (from the original study) with these 109, the total number of experimentally validated read-throughs in Qin *et al* real dataset becomes 144. RTCpredictor re-identified 96 out of 144 read-throughs followed by JAFFA (72) and pizzly (34) while Arriba (2) and InFusion (5) re-identified the lowest number of read-throughs. Moreover, F-measure of RTCpredictor improved and it acquired 2^nd^ spot (Supplementary Table S12).

### Computational requirements

Using one core, RTCpredictor took ~55 mins to complete the run for SRR1657556, but used very high RAM of 164 GB (Figure 5a and Supplementary Table S13). This is because ripgrep software used by RTCpredictor processes all ~nine million junction sequences at once and thereby increasing RAM usage. Although research institutions equipped with high performance computing systems can afford high RAM, it may be challenging for other research institutions with limited resources. Therefore, we introduced a parameter in RTCpredictor which asks users the RAM (in GB) they can comfortably provide. If they give less than 164 GB RAM, RTCpredictor splits junction sequences in multiple parts accordingly and processes each part one by one, thereby reducing RAM usage at the cost of running time (Figure 5a and Supplementary Table S13). The minimum RAM which should be provided to RTCpredictor is 5 GB which is readily available even in present day laptops. Using one core and ~5 GB RAM, RTCpredictor took 109 mins to complete (Figure 5b and Supplementary Table S13). It is still much better than second best method JAFFA which took 525 mins to complete using one core and 5 GB RAM (29). To ramp up the speed of RTCpredictor, we also implemented multiprocessing options using ParallelForkManager perl module where users can use more than one core. With eight cores, RTCpredictor took <20 mins to complete (Figure 5b and Supplementary Table S13).

**Figure 5.**
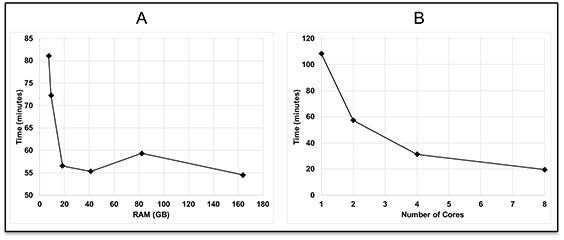
Computational requirements (time and memory) of RTCpredictor when (a) a single core is used, (b) multiple cores are used.

## DISCUSSION

In the quest of chimeric RNAs that have biomarker or therapeutic potentials, read-through class of chimeric RNAs are neglected since their mechanism of formation does not involves chromosomal rearrangement which is one of the classic features of cancer. Therefore, many chimeric RNA prediction methods were designed to purposely discard read-throughs considering them as false positive events. Consequently, the performance of popular chimeric RNA prediction methods on read-through class of chimeric RNAs is poor, especially on real dataset, as demonstrated by the latest benchmark study (29). In this study, we have developed RTCpredictor method specifically for the prediction of read-throughs. We compared RTCpredictor with other methods on read-through datasets and achieved top performance on both simulated and real datasets with a difference of 20% higher sensitivity compared with second-best method on Qin *et al* real dataset. Consistently, RTCpredictor also re-identifying more known read-throughs than other methods on TCGA bladder and colorectal cancer real datasets.

To identify read-throughs, RTCpredictor uses ripgrep to search for ~nine million junction sequences of potential read-throughs against RNA-Seq data. Other chimeric RNA prediction methods use mapping approach and fetch spanning and split reads (also known as junction reads) from mapping step using an aligner and require at least one split/junction read to predict any chimeric RNA. However, as reported by Panagopoulos *et al* (55), a simple grep utility (an in-built command in linux OS), used to search the sequence of *CIC-DUX4* chimeric RNA, outperformed three chimeric RNA prediction methods which used mapping approach. Although grep can be used to search few junction sequences, it cannot handle ~nine million junction sequences. A similar tool ‘agrep’ (56) also allows errors in the match, is faster than grep and has been used for *in silico* validation of chimeric RNAs (37, 57) but it also cannot handle millions of sequences. Like grep, ripgrep performs exact match. It can perform millions of searches in a short time. To accommodate for sequence variations among different individuals, we mapped SNP data on the junction sequences, and added the junction sequence variants in the searchable database.

From a pool of 208 predicted read-throughs obtained from all the software methods on Qin *et al* dataset, 109 were experimentally validated. Here, RTCpredictor outperformed all the other methods and re-identified highest number of experimentally validated read-throughs out of 109. After the addition of 109 and 35 (from original study) read-throughs, the total number of experimentally validated read-throughs increased to 144. As the total number of true positive read-throughs changed, the performance of the methods (Sensitivity, PPV, F-measure) also changed with RTCpredictor retaining the highest sensitivity and moving up three spots in F-measure. In any real dataset, the total number of true positive events is not completely known and there might be more true positive events than the predicted list. Therefore, performance parameters like PPV (fraction of true chimeric RNAs out of total predictions) and F-measure (balance of sensitivity and PPV), are not appropriate for benchmarking chimeric RNA prediction methods, since a tool is penalized if it correctly predicts unknown but true chimeric RNAs. In the latest benchmark study, we observed the same thing on Edgren dataset where originally there were 27 experimentally validated chimeric RNAs which later expanded to 99 resulting in changes in the performance of the methods and thereby their rankings (23, 28, 29, 58).

In terms of computational requirements, using a single core and on the same sample (29), RTCpredictor (~5GB RAM) is 4.8 times faster than the second-best method JAFFA (~5GB RAM) and 6.7 times faster than third best method SOAPfuse (~6GB RAM).

In summary, our newly developed tool RTCpredcitor has the following advantages and some limitations. Advantages of RTCpredictor include: (i) high sensitivity, (ii) fast run time with multiprocessing option, (iii) very easy installation, (iv) can be run on desktop/laptop with 5 GB RAM, (v) can be used on short read data (such as Illumina) or long read data (such as PacBio), (vi) can be used to predict read-throughs from RNA-Seq data of any organism of interest, (vii) compatible with fastq file formats in compressed (*.gz) or uncompressed form, and (viii) provides full sequences of the participating exons for primer designs for experimental validation step.

Limitations of RTCpredictor are: (i) it cannot predict read-throughs whose breakpoints do not map to the end/start of exons and (ii) it may miss few read-throughs, if long reads from Oxford Nanopore platform are used, since the data may have high error rate.

In conclusion, we have developed the first read-through chimeric RNA specific prediction method which outperformed all other methods currently available for chimeric RNA prediction on both simulated and real dataset. We demonstrated its utility by performing experimental testing and validation of 109 read-throughs out of which RTCpredictor re-identified the highest number of read-throughs. We envision that RTCpredictor will be useful in mining the large RNA sequencing datasets and extracting read-throughs with biological/clinical implications, which are otherwise missed by the current methods.

## DECLARATIONS

### Ethics approval and consent to participate

Not applicable.

### Consent for publication

Not applicable.

### Availability of data and materials

All data generated or analyzed during this study are included in this published article. Standalone version of RTCpredictor is available at https://github.com/sandybioteck/RTCpredictor

### Authors’ contributions

S.S. and H.L. conceived the idea. S.S. and S.B.A performed bioinformatic experiments. X.S., S.B.A., T.M., Claire P., Christopher P., Y.S., S.L., and A.S., performed experimental validation of read-throughs. S.S. and H.L. interpreted the data. S.S., X.S. and H.L. wrote the manuscript.

## Acknowledgements

We thank Data Science Institute and the other Computation and Data Resource Exchange (CADRE) partner organizations at the University of Virginia for providing High Performance Computing systems. Author S.S. thank the software development team of Talon Voice software which provides an alternative to type and code using voice commands. All typing work (including manuscript writing and all bioinformatics work) done by author S.S. was done using Talon Voice software which is freely available for Windows, Linux and macOS systems at (https://talonvoice.com/).

## REFERENCES

1. Greger,L., Su,J., Rung,J., Ferreira,P.G., Geuvadis consortium, Lappalainen,T., Dermitzakis,E.T. and Brazma,A. (2014) Tandem RNA chimeras contribute to transcriptome diversity in human population and are associated with intronic genetic variants. PLoS One, 9, e104567.

2. Parra,G., Reymond,A., Dabbouseh,N., Dermitzakis,E.T., Castelo,R., Thomson,T.M., Antonarakis,S.E. and Guigó,R. (2006) Tandem chimerism as a means to increase protein complexity in the human genome. Genome Res, 16, 37–44.

3. Tate,J.G., Bamford,S., Jubb,H.C., Sondka,Z., Beare,D.M., Bindal,N., Boutselakis,H., Cole,C.G., Creatore,C., Dawson,E., et al. (2019) COSMIC: the Catalogue Of Somatic Mutations In Cancer. Nucleic Acids Res, 47, D941–D947.

4. Chwalenia,K., Facemire,L. and Li,H. (2017) Chimeric RNAs in cancer and normal physiology. Wiley Interdiscip Rev RNA, 8.

5. Egashira,S., Jinnin,M., Makino,K., Ajino,M., Shimozono,N., Okamoto,S., Tazaki,Y., Hirano,A., Ide,M., Kajihara,I., et al. (2019) Recurrent Fusion Gene ADCK4-NUMBL in Cutaneous Squamous Cell Carcinoma Mediates Cell Proliferation. J Invest Dermatol, 139, 954–957.

6. Wu,H., Li,X. and Li,H. (2019) Gene fusions and chimeric RNAs, and their implications in cancer. Genes Dis, 6, 385–390.

7. Finta,C. and Zaphiropoulos,P.G. (2002) Intergenic mRNA molecules resulting from trans-splicing. J Biol Chem, 277, 5882–90.

8. Wang,L., Xiong,X., Yao,Z., Zhu,J., Lin,Y., Lin,W., Li,K., Xu,X., Guo,Y., Chen,Y., et al. (2021) Chimeric RNA ASTN2-PAPPAas aggravates tumor progression and metastasis in human esophageal cancer. Cancer Lett, 501, 1–11.

9. Rowley,J.D. (1973) Letter: A new consistent chromosomal abnormality in chronic myelogenous leukaemia identified by quinacrine fluorescence and Giemsa staining. Nature, 243, 290–3.

10. Tomlins,S.A., Laxman,B., Varambally,S., Cao,X., Yu,J., Helgeson,B.E., Cao,Q., Prensner,J.R., Rubin,M.A., Shah,R.B., et al. (2008) Role of the TMPRSS2-ERG gene fusion in prostate cancer. Neoplasia, 10, 177–88.

11. Linardic,C.M. (2008) PAX3-FOXO1 fusion gene in rhabdomyosarcoma. Cancer Lett, 270, 10–8.

12. Lin,Y., Dong,H., Deng,W., Lin,W., Li,K., Xiong,X., Guo,Y., Zhou,F., Ma,C., Chen,Y., et al. (2019) Evaluation of Salivary Exosomal Chimeric GOLM1-NAA35 RNA as a Potential Biomarker in Esophageal Carcinoma. Clin Cancer Res, 25, 3035–3045.

13. Kannan,K., Wang,L., Wang,J., Ittmann,M.M., Li,W. and Yen,L. (2011) Recurrent chimeric RNAs enriched in human prostate cancer identified by deep sequencing. Proc Natl Acad Sci U S A, 108, 9172–7.

14. Zhou,J., Liao,J., Zheng,X. and Shen,H. (2012) Chimeric RNAs as potential biomarkers for tumor diagnosis. BMB Rep, 45, 133–40.

15. Babiceanu,M., Qin,F., Xie,Z., Jia,Y., Lopez,K., Janus,N., Facemire,L., Kumar,S., Pang,Y., Qi,Y., et al. (2016) Recurrent chimeric fusion RNAs in non-cancer tissues and cells. Nucleic Acids Res, 44, 2859–72.

16. Singh,S., Qin,F., Kumar,S., Elfman,J., Lin,E., Pham,L.-P., Yang,A. and Li,H. (2020) The landscape of chimeric RNAs in non-diseased tissues and cells. Nucleic Acids Res, 48, 1764–1778.

17. Mukherjee,S., Detroja,R., Balamurali,D., Matveishina,E., Medvedeva,Y.A., Valencia,A., Gorohovski,A. and Frenkel-Morgenstern,M. (2021) Computational analysis of sense-antisense chimeric transcripts reveals their potential regulatory features and the landscape of expression in human cells. NAR Genom Bioinform, 3, lqab074.

18. Novo,F.J., de Mendíbil,I.O. and Vizmanos,J.L. (2007) TICdb: a collection of gene-mapped translocation breakpoints in cancer. BMC Genomics, 8, 33.

19. Balamurali,D., Gorohovski,A., Detroja,R., Palande,V., Raviv-Shay,D. and Frenkel-Morgenstern,M. (2019) ChiTaRS 5.0: the comprehensive database of chimeric transcripts matched with druggable fusions and 3D chromatin maps. Nucleic Acids Res, 10.1093/nar/gkz1025.

20. Hu,X., Wang,Q., Tang,M., Barthel,F., Amin,S., Yoshihara,K., Lang,F.M., Martinez-Ledesma,E., Lee,S.H., Zheng,S., et al. (2018) TumorFusions: an integrative resource for cancer-associated transcript fusions. Nucleic Acids Res, 46, D1144–D1149.

21. Jang,Y.E., Jang,I., Kim,S., Cho,S., Kim,D., Kim,K., Kim,J., Hwang,J., Kim,S., Kim,J., et al. (2020) ChimerDB 4.0: an updated and expanded database of fusion genes. Nucleic Acids Res, 48, D817–D824.

22. Akiva,P., Toporik,A., Edelheit,S., Peretz,Y., Diber,A., Shemesh,R., Novik,A. and Sorek,R. (2006) Transcription-mediated gene fusion in the human genome. Genome Res, 16, 30–6.

23. Davidson,N.M., Majewski,I.J. and Oshlack,A. (2015) JAFFA: High sensitivity transcriptome-focused fusion gene detection. Genome Med, 7, 43.

24. Denoeud,F., Kapranov,P., Ucla,C., Frankish,A., Castelo,R., Drenkow,J., Lagarde,J., Alioto,T., Manzano,C., Chrast,J., et al. (2007) Prominent use of distal 5’ transcription start sites and discovery of a large number of additional exons in ENCODE regions. Genome Res, 17, 746–59.

25. Li,Y., Heavican,T.B., Vellichirammal,N.N., Iqbal,J. and Guda,C. (2017) ChimeRScope: a novel alignment-free algorithm for fusion transcript prediction using paired-end RNA-Seq data. Nucleic Acids Res, 45, e120.

26. Uhrig,S., Fröhlich,M., Hutter,B. and Brors,B. (2018) PO-400 Arriba – fast and accurate gene fusion detection from RNA-seq data. ESMO Open, 3, A179 LP–A179.

27. Uhrig,S., Ellermann,J., Walther,T., Burkhardt,P., Fröhlich,M., Hutter,B., Toprak,U.H., Neumann,O., Stenzinger,A., Scholl,C., et al. (2021) Accurate and efficient detection of gene fusions from RNA sequencing data. Genome Res, 31, 448–460.

28. Edgren,H., Murumagi,A., Kangaspeska,S., Nicorici,D., Hongisto,V., Kleivi,K., Rye,I.H., Nyberg,S., Wolf,M., Borresen-Dale,A.-L., et al. (2011) Identification of fusion genes in breast cancer by paired-end RNA-sequencing. Genome Biol, 12, R6.

29. Singh,S. and Li,H. (2021) Comparative study of bioinformatic tools for the identification of chimeric RNAs from RNA Sequencing. RNA Biol, 18, 254–267.

30. Rickman,D.S., Pflueger,D., Moss,B., VanDoren,V.E., Chen,C.X., de la Taille,A., Kuefer,R., Tewari,A.K., Setlur,S.R., Demichelis,F., et al. (2009) SLC45A3-ELK4 is a novel and frequent erythroblast transformation-specific fusion transcript in prostate cancer. Cancer Res, 69, 2734–8.

31. Zhang,Y., Gong,M., Yuan,H., Park,H.G., Frierson,H.F. and Li,H. (2012) Chimeric transcript generated by cis-splicing of adjacent genes regulates prostate cancer cell proliferation. Cancer Discov, 2, 598–607.

32. Varley,K.E., Gertz,J., Roberts,B.S., Davis,N.S., Bowling,K.M., Kirby,M.K., Nesmith,A.S., Oliver,P.G., Grizzle,W.E., Forero,A., et al. (2014) Recurrent read-through fusion transcripts in breast cancer. Breast Cancer Res Treat, 146, 287–97.

33. Nacu,S., Yuan,W., Kan,Z., Bhatt,D., Rivers,C.S., Stinson,J., Peters,B.A., Modrusan,Z., Jung,K., Seshagiri,S., et al. (2011) Deep RNA sequencing analysis of readthrough gene fusions in human prostate adenocarcinoma and reference samples. BMC Med Genomics, 4, 11.

34. Wu,P., Yang,S., Singh,S., Qin,F., Kumar,S., Wang,L., Ma,D. and Li,H. (2018) The Landscape and Implications of Chimeric RNAs in Cervical Cancer. EBioMedicine, 37, 158–167.

35. Qin,F., Song,Z., Chang,M., Song,Y., Frierson,H. and Li,H. (2016) Recurrent cis-SAGe chimeric RNA, D2HGDH-GAL3ST2, in prostate cancer. Cancer Lett, 380, 39–46.

36. Zhu,D., Singh,S., Chen,X., Zheng,Z., Huang,J., Lin,T. and Li,H. (2019) The landscape of chimeric RNAs in bladder urothelial carcinoma. Int J Biochem Cell Biol, 110, 50–58.

37. Wu,H., Singh,S., Xie,Z., Li,X. and Li,H. (2020) Landscape characterization of chimeric RNAs in colorectal cancer. Cancer Lett, 489, 56–65.

38. Tang,Y., Qin,F., Liu,A. and Li,H. (2017) Recurrent fusion RNA DUS4L-BCAP29 in non-cancer human tissues and cells. Oncotarget, 8, 31415–31423.

39. Zhuo,J.-S., Jing,X.-Y., Du,X. and Yang,X.-Q. (2018) Generation of Chimeric RNAs by cis-splicing of adjacent genes (cis-SAGe) in mammals. Yi Chuan, 40, 145–154.

40. Elfman,J. and Li,H. (2018) Chimeric RNA in Cancer and Stem Cell Differentiation. Stem Cells Int, 2018, 3178789.

41. Rodríguez-Martín,B., Palumbo,E., Marco-Sola,S., Griebel,T., Ribeca,P., Alonso,G., Rastrojo,A., Aguado,B., Guigó,R. and Djebali,S. (2017) ChimPipe: accurate detection of fusion genes and transcription-induced chimeras from RNA-seq data. BMC Genomics, 18, 7.

42. Qin,F., Song,Z., Babiceanu,M., Song,Y., Facemire,L., Singh,R., Adli,M. and Li,H. (2015) Discovery of CTCF-sensitive Cis-spliced fusion RNAs between adjacent genes in human prostate cells. PLoS Genet, 11, e1005001.

43. Patel,R.K. and Jain,M. (2012) NGS QC Toolkit: a toolkit for quality control of next generation sequencing data. PLoS One, 7, e30619.

44. Zhu,D., Singh,S., Chen,X., Zheng,Z., Huang,J., Lin,T. and Li,H. (2019) The landscape of chimeric RNAs in bladder urothelial carcinoma. Int J Biochem Cell Biol, 110, 50–58.

45. Liu,S., Tsai,W.-H., Ding,Y., Chen,R., Fang,Z., Huo,Z., Kim,S., Ma,T., Chang,T.-Y., Priedigkeit,N.M., et al. (2016) Comprehensive evaluation of fusion transcript detection algorithms and a meta-caller to combine top performing methods in paired-end RNA-seq data. Nucleic Acids Res, 44, e47.

46. Qin,F., Song,Y., Zhang,Y., Facemire,L., Frierson,H. and Li,H. (2016) Role of CTCF in Regulating SLC45A3-ELK4 Chimeric RNA. PLoS One, 11, e0150382.

47. Quinlan,A.R. and Hall,I.M. (2010) BEDTools: a flexible suite of utilities for comparing genomic features. Bioinformatics, 26, 841–2.

48. Jia,W., Qiu,K., He,M., Song,P., Zhou,Q., Zhou,F., Yu,Y., Zhu,D., Nickerson,M.L., Wan,S., et al. (2013) SOAPfuse: an algorithm for identifying fusion transcripts from paired-end RNA-Seq data. Genome Biol, 14, R12.

49. Benelli,M., Pescucci,C., Marseglia,G., Severgnini,M., Torricelli,F. and Magi,A. (2012) Discovering chimeric transcripts in paired-end RNA-seq data by using EricScript. Bioinformatics, 28, 3232–9.

50. Nicorici,D., Şatalan,M., Edgren,H., Kangaspeska,S., Murumägi,A., Kallioniemi,O., Virtanen,S. and Kilkku,O. (2014) FusionCatcher – a tool for finding somatic fusion genes in paired-end RNA-sequencing data. bioRxiv, 10.1101/011650.

51. Melsted,P., Hateley,S., Joseph,I.C., Pimentel,H., Bray,N. and Pachter,L. (2017) Fusion detection and quantification by pseudoalignment. bioRxiv, 10.1101/166322.

52. Okonechnikov,K., Imai-Matsushima,A., Paul,L., Seitz,A., Meyer,T.F. and Garcia-Alcalde,F. (2016) InFusion: Advancing Discovery of Fusion Genes and Chimeric Transcripts from Deep RNA-Sequencing Data. PLoS One, 11, e0167417.

53. Vu,T.N., Deng,W., Trac,Q.T., Calza,S., Hwang,W. and Pawitan,Y. (2018) A fast detection of fusion genes from paired-end RNA-seq data. BMC Genomics, 19, 786.

54. Iyer,M.K., Chinnaiyan,A.M. and Maher,C.A. (2011) ChimeraScan: a tool for identifying chimeric transcription in sequencing data. Bioinformatics, 27, 2903–4.

55. Panagopoulos,I., Gorunova,L., Bjerkehagen,B. and Heim,S. (2014) The ‘grep’ command but not FusionMap, FusionFinder or ChimeraScan captures the CIC-DUX4 fusion gene from whole transcriptome sequencing data on a small round cell tumor with t(4;19)(q35;q13). PLoS One, 9, e99439.

56. Wu,S. (University of A. and Manber,U. (University of A. (1990) AGREP - A Fast Approximate Pattern-matching Tool. In Proceedings of the Winter 1990 USENIX Conference. San Francisco, pp. 153–162.

57. Singh,S. and Li,H. (2020) Prediction, Characterization, and In Silico Validation of Chimeric RNAs. Methods Mol Biol, 2079, 3–12.

58. Kangaspeska,S., Hultsch,S., Edgren,H., Nicorici,D., Murumägi,A. and Kallioniemi,O. (2012) Reanalysis of RNA-sequencing data reveals several additional fusion genes with multiple isoforms. PLoS One, 7, e48745.

