## Supplementary figures and images for "RTCpredictor: Identification of Read-Through Chimeric RNAs from RNA Sequencing Data"

### Fig S1

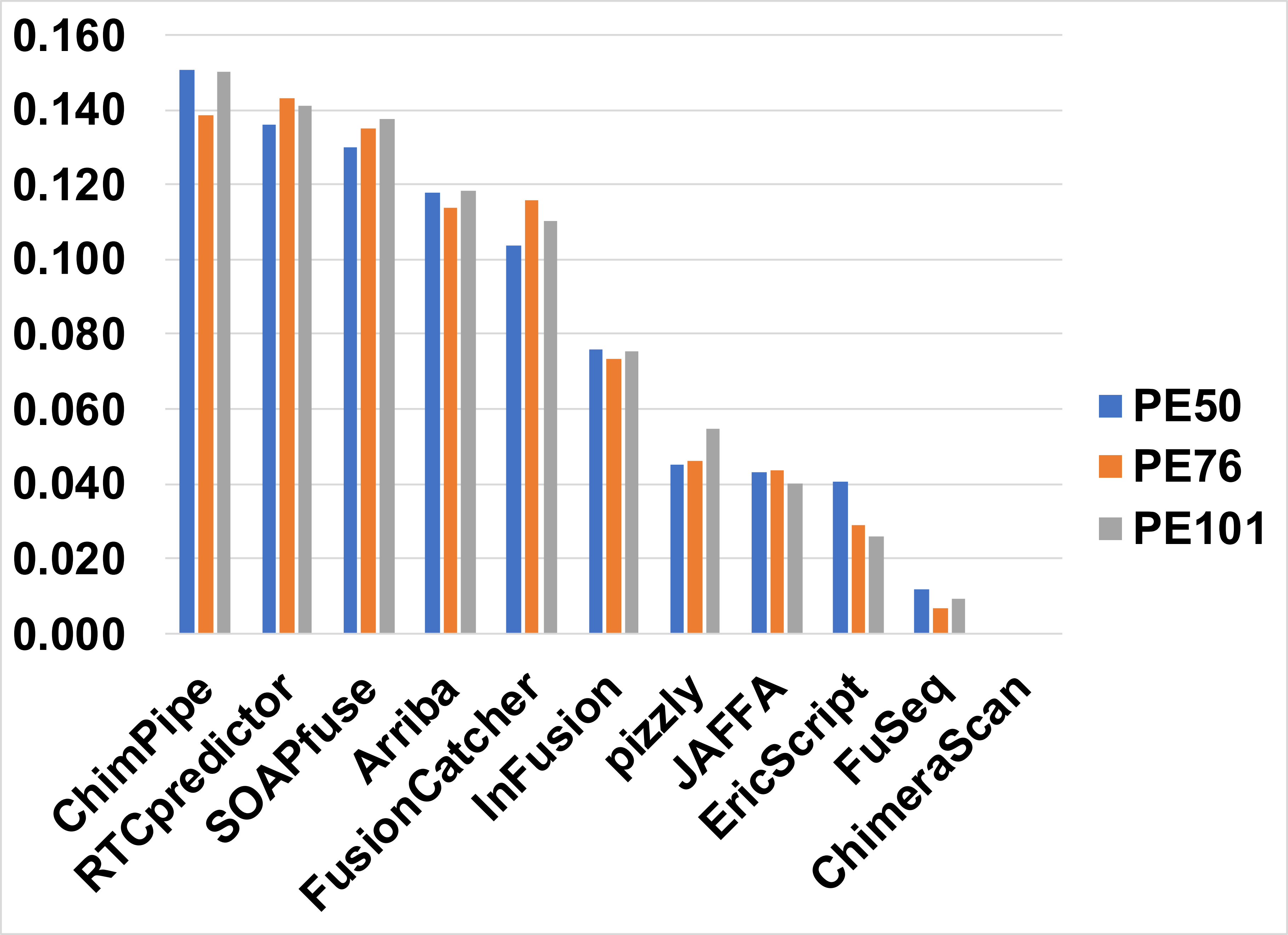

### Fig S2

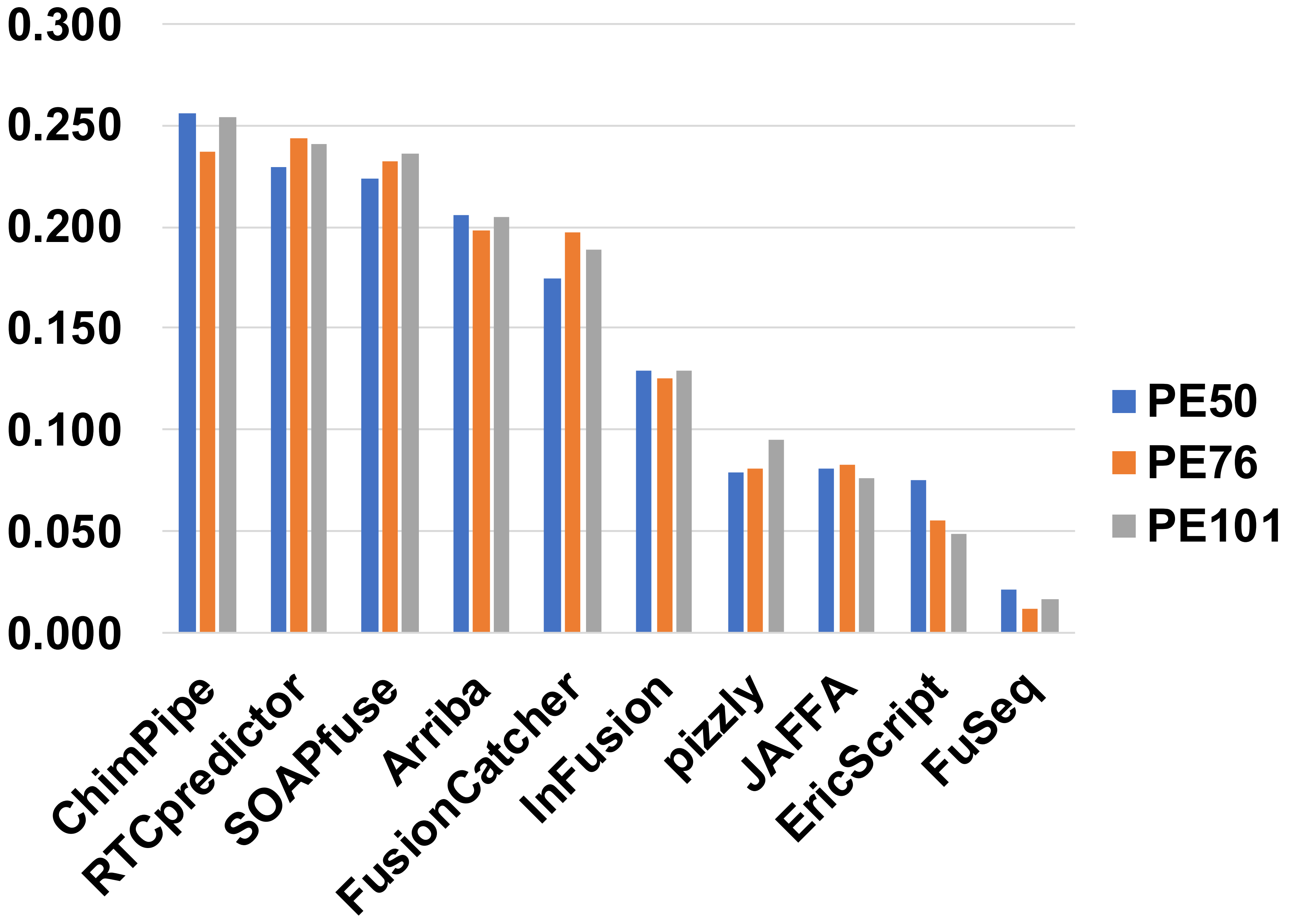

### Fig S3

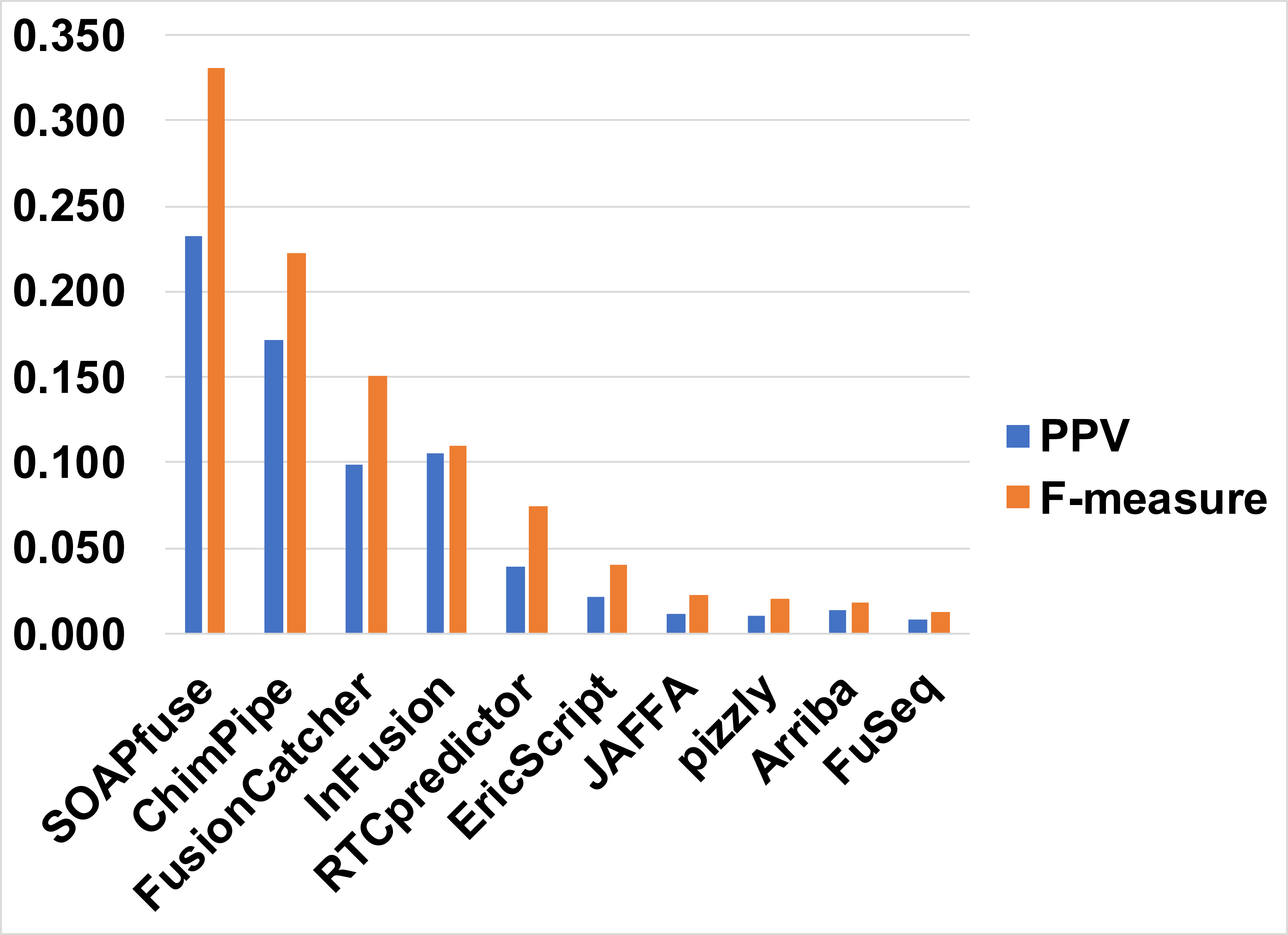
